# Temporal Interference Modulates Medial Temporal and Frontoparietal Activity During Mnemonic Discrimination: A high-resolution whole-brain fMRI investigation

**DOI:** 10.64898/2026.03.03.708601

**Authors:** Min Sung Seo, Rebecca L. Wagner, Jordan D. Chamberlain, Nancy A. Dennis

## Abstract

Mnemonic discrimination—the ability to distinguish between existing memories and similar new inputs—supports accurate memory in the face of interference (e.g., remembering where you parked today versus yesterday in the same parking deck). Although hippocampal pattern separation has been emphasized as a key mechanism, recent theoretical frameworks and empirical findings suggest that surrounding medial temporal lobe and cortical regions also contribute by resolving interference. An important source of interference in everyday life is temporal interference: similar events may compete not only because they overlap perceptually, but also because they are separated by intervening experiences (e.g., having to remember your morning parking spot after a full day of work). Here, we investigated how temporal interference impacts mnemonic discrimination and its neural correlates using high-resolution fMRI and a continuous recognition task in which the lag between an item’s first presentation and its subsequent target or similar lure was systematically manipulated (10, 60, or 140 intervening trials). Behaviorally, increasing lag impaired both mnemonic discrimination and target recognition. At the neural level, lag effects were not observed in hippocampal subfields. Rather, temporal interference modulated activity in extrahippocampal MTL regions including parahippocampal and perirhinal cortex, as well as distributed cortical regions including frontoparietal control areas. Importantly, lag-related modulation was process-specific, with partially dissociable patterns for successful mnemonic discrimination and target recognition. Together, these findings demonstrate that temporal interference engages a distributed MTL-cortical network that supports interference resolution during mnemonic discrimination.

## Introduction

An important aspect of episodic memory is mnemonic discrimination: the ability to distinguish between existing memories and similar, yet new inputs (e.g., remembering where you parked today versus yesterday in the same parking deck). Neurocomputational models propose that successful mnemonic discrimination depends on pattern separation, a process that transforms overlapping inputs into distinct neural representations, thereby reducing competition among similar memories (McClelland et al., 1995; Norman & O’Reilly, 2003; O’Reilly & Rudy, 2001; Sahay et al., 2011). Importantly, converging evidence from multiple methodologies including lesion studies (Kirwan et al., 2012; Reyes et al., 2018; Schmidt et al., 2012), animal research (McAllister et al., 2013), computational modeling (McClelland et al., 2020; O’Reilly & Rudy, 2001; Treves & Rolls, 1994), and human neuroimaging studies (Bakker et al., 2008; Kirwan & Stark, 2007), point to the hippocampus as a critical brain structure that supports pattern separation and mnemonic discrimination.

To investigate pattern separation in humans, a large body of work has employed mnemonic discrimination tasks (MDT). In these tasks, participants typically view images of everyday objects during encoding, and later engage in a recognition test where they view exact repetitions of studied items (targets), similar but non-identical items (lures), and entirely novel items (foils). During this recognition test, participants must identify images as being the “same” as previously viewed images, “similar” to previously viewed images, or entirely “new.” Importantly, endorsing lures as “similar” instead of “same” on MDTs is taken as a behavioral index of mnemonic discrimination, as participants must engage in pattern separation to accurately dissociate test lures from studied targets. Neurally, functional magnetic resonance imaging (fMRI) studies using MDTs have shown that mnemonic discrimination performance is selectively sensitive to hippocampal integrity (both functional and structural), in contrast to simple object recognition performances (for reviews, see Leal & Yassa, 2018; Stark et al., 2019).

Early fMRI work using MDTs demonstrated that hippocampal activity differentiates successful lure correct rejections (correctly endorsing lures as being “similar” to studied targets) from lure false alarms (erroneously endorsing lures as being the “same” as studied targets), whereas other medial temporal lobe (MTL) regions respond similarly to correct target-same and lure-similar responses, suggesting that hippocampal signals uniquely reflect the resolution of mnemonic overlap rather than retrieval success per se (Kirwan & Stark, 2007). Subsequent studies extended these findings by implicating specific hippocampal subfields, particularly the dentate gyrus (DG) and CA3, in successful pattern separation. For example, Bakker et al. (2008) reported that CA3/DG activity showed a strong bias toward pattern separation, exhibiting unique responses to lures relative to exact target repetitions, whereas other hippocampal and MTL regions did not show this differentiation. Furthermore, 7T fMRI studies have demonstrated greater DG engagement during successful lure discrimination compared to target repetition trials (Berron et al., 2016; Yang et al., 2024). Collectively, fMRI studies using MDTs support the view that mnemonic discrimination depends critically on hippocampal computations within the CA3 and particularly the DG, that transform overlapping inputs into distinct mnemonic representations.

While there are various components to the MDT, a key aspect for evaluating pattern separation is mnemonic interference. To tax hippocampal pattern separation processes, lures are constructed to overlap with studied targets, creating competition between similar memory representations at retrieval. In many MDTs, this competition creates a source of interference that is manipulated via perceptual similarity (i.e., lure similarity). Lure items may vary along a continuum from highly similar, to minimally similar, relative to their corresponding studied items. Critically, as lure similarity increases, interference also increases, and successful mnemonic discrimination becomes more difficult. Increasing interference places greater demands on hippocampal pattern separation mechanisms, such that small differences in input must be transformed into disproportionately larger differences in mnemonic representations (Guzowski et al., 2004; Yassa & Stark, 2011). Consistent with this account, behavioral studies show that mnemonic discrimination performance declines as lure similarity increases across various populations (healthy younger and older adults, and cognitively impaired older adults; Holden et al., 2012; Reagh & Yassa, 2014; Stark et al., 2010) and different stimulus features such as visual similarity and spatial proximity (Holden et al., 2012; Motley & Kirwan, 2012; Reagh et al., 2014).

Another important and ecologically relevant source of interference is the temporal distance between similar episodic events (i.e., temporal interference). In everyday memory, interference often arises not only because two events are perceptually similar, but also because similar events can be separated in time (e.g., having to remember your morning parking spot after a full day of work). Greater temporal separation increases the likelihood that overlapping episodic representations compete during retrieval. In most MDT variants, encoding and retrieval are divided into separate phases. During encoding, participants typically engage in an incidental encoding task while viewing stimuli (e.g., indoor/outdoor judgments for pictures of everyday objects). Subsequently, participants complete a recognition memory test in which they make judgments to targets, lures, and foils. Using this format, studies have manipulated the delay between encoding and retrieval (e.g., immediate vs. one-week delay) and have shown that mnemonic discrimination performance declines with longer delays (Leal et al., 2014, 2019; Morales-Calva et al., 2026).

Although delayed versus immediate retrieval comparisons provide useful insights into how interference associated with retrieval delay impacts mnemonic discrimination, they do not allow systematic manipulation of the temporal distance between individual pairs of related stimuli (i.e., a specific item and its later lure). In contrast, continuous-recognition versions of the MDT allow stimuli to be presented within a single stream, enabling more precise control over the number of intervening items, and timing, between an initial presentation and its subsequent target or lure (i.e., lag). This design affords a more fine-grained investigation of temporal interference at the level of individual mnemonic representations. Similar to lure similarity effects, prior behavioral work has shown that increasing lag leads to reduced mnemonic discrimination performance (Roberts et al., 2014; S. M. Stark et al., 2015; Tolentino et al., 2012). Based on interference-based models of hippocampal memory, greater lag should induce more temporal interference, resulting in increased demands on hippocampal pattern separation mechanisms to support successful mnemonic discrimination (Kumaran et al., 2016; McClelland et al., 1995; Norman & O’Reilly, 2003; Yassa & Reagh, 2013). However, despite theoretical predictions and robust behavioral lag effects, relatively little is known about how temporal interference affects the neural mechanisms supporting mnemonic discrimination.

While much of the human pattern separation literature has focused on the hippocampus and its subfields, accumulating theoretical and empirical work challenges a primarily hippocampus-centric view. The Cortico-Hippocampal Pattern Separation (CHiPS) framework (Amer & Davachi, 2023) proposes that pattern separation emerges from coordinated interactions across sensory cortices, frontoparietal regions, and medial temporal lobe (MTL) structures. According to CHiPS, extrahippocampal regions contribute to pattern separation via: (1) interference resolution in sensory/perceptual cortices reducing representational overlap before information reaches the hippocampus; and (2) top-down modulation from cognitive control regions biasing hippocampal computations toward greater differentiation when task demands require fine-grained discrimination. In this view, pattern separation is not solely an intrinsic hippocampal computation, but rather, the outcome of cortical feature filtering, domain-specific MTL processing, and hippocampal amplification of pre-differentiated inputs. The CHiPS model thus predicts that neural signals related to pattern separation should be detectable not only in hippocampal subfields, but also in cortical regions that resolve interference and shape the inputs received by the hippocampus (Amer & Davachi, 2023).

Consistent with the CHiPS framework, several fMRI studies have reported extrahippocampal contributions to pattern separation. For example, Paleja et al. (2014) demonstrated that successful lure discrimination engages a distributed hippocampal-cortical network spanning occipitotemporal regions, lateral prefrontal cortex, parietal cortex, and cerebellum, with functional connectivity between the hippocampus and these regions predicting successful mnemonic discrimination. Similarly, Pidgeon and Morcom (2016) showed that perceptual regions in occipitotemporal cortex encode item-specific features that support mnemonic discrimination, while Wais et al. (2017) reported that successful mnemonic discrimination was associated with strengthened connectivity between MTL regions (e.g., entorhinal cortex, hippocampus), and frontal, parietal cortices (e.g., inferior frontal gyrus and angular gyrus). Reagh et al. (2018) also demonstrated that perirhinal (PRC) and lateral entorhinal cortices support feature-level differentiation of similar objects, providing evidence that non-hippocampal MTL regions directly contribute to pattern separation.

More recent high-resolution fMRI studies have provided converging evidence for the interaction of hippocampal-cortical regions supporting pattern separation processes. For instance, Klippenstein et al. (2020) reported that the blood-oxygen-level-dependent (BOLD) activity in the hippocampus and occipital regions was associated with pattern separation, and Stevenson et al. (2020) found that greater activity in CA3/DG and perirhinal cortex was correlated with successful mnemonic discrimination. Using multiband 3T fMRI (Xu et al., 2013), Nash et al. (2021) were able to isolate hippocampal subfields more precisely and identified pattern separation-related neural signals in the subiculum as well as in cortical regions such as the parahippocampal, prefrontal, and parietal cortices. Most recently, Yang et al. (2024) employed 7T whole-brain high-resolution fMRI and identified neural correlates of successful mnemonic discrimination in frontoparietal, temporal, and subcortical regions, alongside CA3/DG within the hippocampus, emphasizing cortico-hippocampal contributions to pattern separation in the MDT. Nevertheless, despite growing evidence for a joint hippocampal-cortical contribution to pattern separation, it remains unclear how temporal interference modulates these cortico-hippocampal interactions.

The current study employed high-resolution whole-brain fMRI to investigate the effects of temporal interference on mnemonic discrimination and its neural correlates in healthy younger adults. Temporal interference was manipulated using a continuous-recognition formatted MDT (S. M. Stark et al., 2019), in which the lag between an item’s initial presentation and its subsequent target or lure was systematically varied across three discrete distances (10, 60, and 140). In contrast to studies in which lag varied continuously across a broad range (e.g., 20–100) and was later binned into categories (e.g., short, medium, long; Kirwan & Stark, 2007; Stark et al., 2015), the present design implemented discrete lag values with equal numbers of trials at each lag. This approach provided greater experimental control over temporal interference, reduced variability within lag conditions, and enabled a more precise investigation of how temporal distance affects mnemonic discrimination. Inside the MRI scanner, participants viewed images of everyday objects that included first presentations (later repeated exactly as targets or shown as similar lures), exact repetitions of previously presented objects (targets), similar but non-identical images of previously presented objects (lures), and new objects presented only once (foil). Participants made same/similar/new judgments on each trial. Critically, temporal interference was manipulated by setting the lag for targets and lures such that the number of intervening trials between an item’s first presentation and its target or lure was 10, 60 or 140.

Based on interference-based models of hippocampal memory and prior MDT studies (Roberts et al., 2014; S. M. Stark et al., 2015, 2019; Tolentino et al., 2012; Yassa & Reagh, 2013), we hypothesized that mnemonic discrimination performance would decline as lag increased, reflecting greater temporal interference effects. At the neural level, we hypothesized that DG activity would increase with lag during successful mnemonic discrimination, as greater interference should place increased demands on pattern separation computations. In addition, consistent with the CHiPS model (Amer & Davachi, 2023), we expected successful mnemonic discrimination to be associated with activity in frontoparietal control regions, as well as ventral visual and MTL regions (perirhinal and entorhinal cortex). Within the CHiPS framework, increasing temporal interference should not only increase hippocampal pattern separation, but also increase the contribution of cortical regions involved in interference resolution. Accordingly, we hypothesized that activity in ventral visual cortex and perirhinal cortex would increase with lag during successful lure discrimination, reflecting greater demands on feature-level and object-level differentiation. We also expected lag-related activity increases in frontoparietal control regions, consistent with greater top-down engagement under increased interference. Critically, these lag effects were predicted to be specific to successful mnemonic discrimination.

## Methods

### Participants

Thirty-four healthy young adults were recruited via physical advertisements and word of mouth at The Pennsylvania State University, University Park campus. All participants were screened for 3T MRI safety and excluded if they reported colorblindness, psychiatric or neurological disorders, a history of head injury or stroke, learning disabilities, substance abuse, or use of any medications affecting cognitive, cardiovascular, or neural function. All participants also reported being right-handed. Of the 34 participants, two were excluded for not completing the entire experiment protocol, two were excluded for failing to follow task instructions, and one was excluded after being found ineligible for MRI scanning following consent. The final sample comprised 29 subjects (20 females; *M_age_* = 21.38, *SD_age_* = 3.18, range*_age_*: 18–30; *M_education_* = 14.76, *SD_education_* = 1.96). All participants provided written informed consent to participate in the study, with all experimental protocols approved by The Pennsylvania State University Institutional Review Board.

### Procedure

All participants completed a continuous variant of the MDT (Stark et al., 2013; Stark et al., 2015), in which they made a memory response for each item without a separate study phase. Prior to entering the MRI, participants were briefed on the continuous nature of the memory task, that they would need to respond to each trial that appeared, and the available response options: “same”, “similar”, and “new”. “Same” responses were to be used when participants believed an image to be the exact same image they had seen earlier in the task. “Similar” responses were to be used when participants believed an image was similar to one they had seen earlier, but the image was not exactly the same as the one they had previously seen. Finally, “new” responses were to be used when they believed an image to be completely different from anything they had seen previously in the task. With these verbal and written instructions, participants were shown example images that did not overlap with any images shown during the memory task and were provided explicit response instruction for each example image.

Upon entering the MRI scanner, participants first underwent standard localizer, scout, anatomical, and hippocampal slab image acquisition scans, which lasted about 10 minutes. They then received a written reminder of the same/similar/new response instructions before beginning the task portion of the scan. The MDT was administered using E-Prime 3.0 (Psychology Software Tools, Pittsburgh, PA, USA) and consisted of two runs. Each run contained 300 images selected from a publicly available object-lure stimulus database (https://github.com/celstark/MST; Stark et al., 2019). Within each run, 120 images were presented as “first-presentation” trials. Later in the trial, 60 of these images were re-presented as targets, and a different 60 new images served as lures that were visually similar (but not identical) to one of the earlier images. All lure images were of the same visual similarity level (lure discriminability level 4; Stark et al., 2019). For both targets and lures, 20 of each were presented 10 trials after its corresponding first presentation, 20 of each were presented after 60 trials, and another 20 after 140 trials. The remaining 60 trials in each run were “foil” images that were never re-presented (see Figure 1 for schematic depiction of the task design). Each image was presented for 2500 ms, followed by a 500 ms interstimulus interval (ISI). Participants made a memory judgment for every image using an MRI-compatible button box with three buttons corresponding to “same”(index finger), “similar”(middle finger), and “new”(ring finger). Before the second run began, participants were reminded of the response instructions. The second run used a different set of images but maintained the same number of first-presentation, target, lure, and foil trials, as well as the same presentation timing and lag structure as the first run. The order of trial presentation (both within and across run) was identical for all participants. In total, each participant viewed and responded to 600 images over approximately 30 minutes in the scanner.

**Figure 1.**
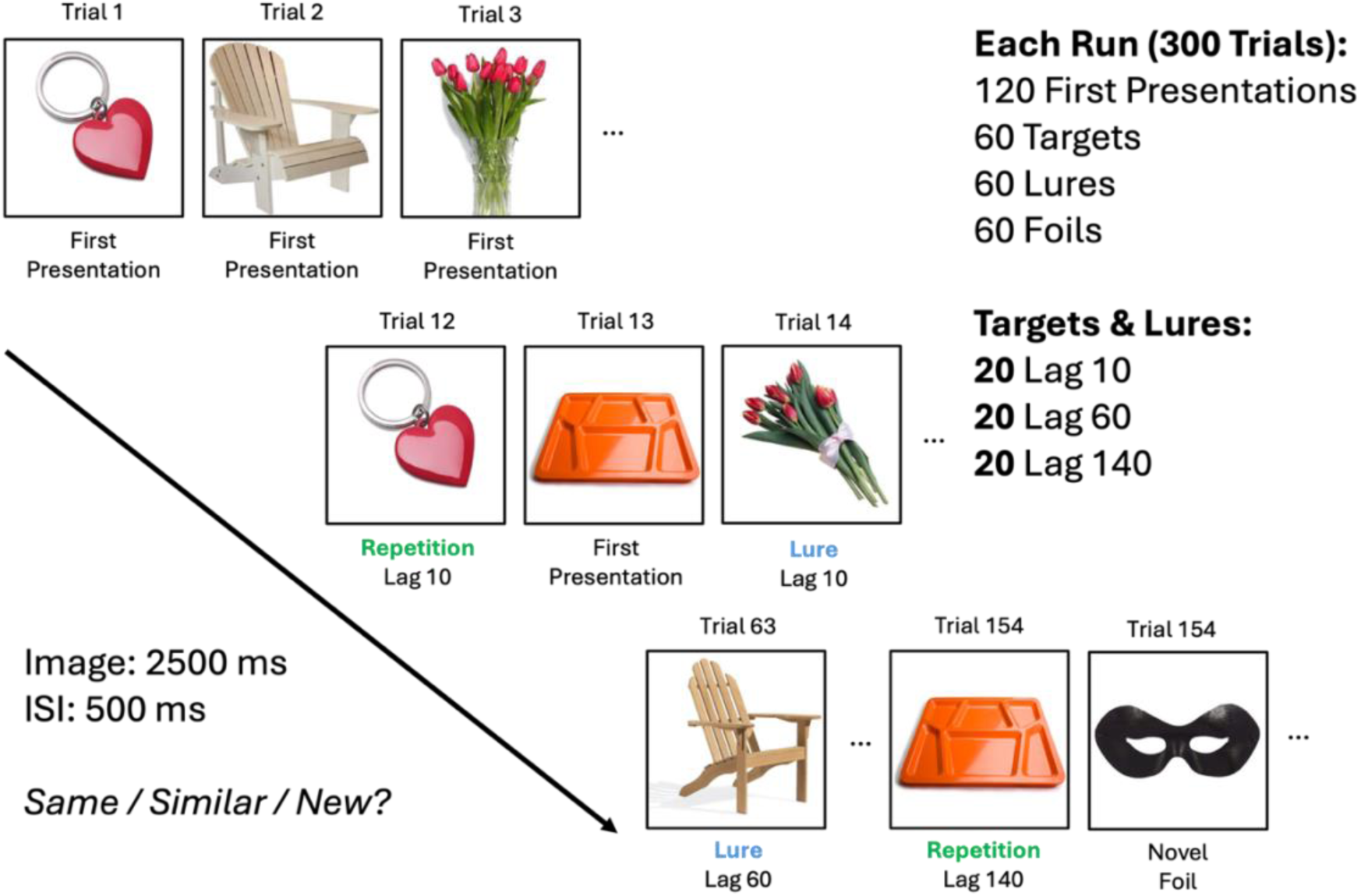
Experimental design schematic. Participants viewed a continuous series of object images (2500 ms stimulus presentation, 500 ms inter-stimulus interval; ISI) and made a memory judgment on each trial (“same,” “similar,” or “new”). Trial types included first presentations (later repeated exactly as targets or presented as similar lures), targets, lures, and foils (novel image presented only once). Targets and lures appeared after their corresponding first presentations at one of three lag intervals (10, 60 and 140 intervening trials). The task consisted of two runs.

### Image acquisition

Structural and functional brain images were acquired using a Siemens 3T scanner equipped with a 64-channel head coil. A T1-weighted sagittal localizer was collected to locate the anterior (AC) and posterior (PC) commissures for anatomical alignment and the slice acquisition boundaries, which were then applied to the anatomical and functional scans. The anatomical image was acquired using a 3D multi-planar reduced angle gradient-echo (MPRAGE) technique with a 1700 ms TR, 2.28 ms TE, 256 mm field of view (FOV), and 192 1mm thick axial slices resulting in 1mm isotropic voxels. Functional images were acquired using interleaved functional echo planar imaging (EPI) with a multiband factor of two, a 2500ms TR, 32.00 ms TE, 221 mm FOV, and 72 1.7mm thick slices resulting in 1.7mm isotropic voxels. To collect hippocampal subfields, an additional high-resolution proton density (pd)-weighted T2 structural slab image was acquired. This slab was auto-aligned perpendicular to the longitudinal axis of the hippocampus and visually confirmed prior to the collection of the image, which had a 6490 ms TR, 16 ms TE, 206 mm FOV, and 33 2mm thick slices, resulting in a voxel size of 0.4 x 0.4 x 2 mm^3^.

### fMRIPrep preprocessing

Results included in this manuscript come from preprocessing performed using fMRIPrep 20.1.1 (Esteban et al., 2019).

#### Anatomical data preprocessing

The T1-weighted (T1w) image was corrected for intensity non-uniformity (INU) with N4BiasFieldCorrection (Tustison et al., 2010), distributed with ANTs 2.2.0 (B. Avants et al., 2008), and used as T1w-reference throughout the workflow. The T1w-reference was then skull-stripped with a Nipype implementation of the antsBrainExtraction.sh workflow (from ANTs), using OASIS30ANTs as target template. Brain tissue segmentation of cerebrospinal fluid (CSF), white-matter (WM) and gray-matter (GM) was performed on the brain-extracted T1w using fast (FSL 5.0.9, Zhang et al., 2001). Brain surfaces were reconstructed using recon-all (FreeSurfer 6.0.1, Dale et al., 1999) and the brain mask estimated previously was refined with a custom variation of the method to reconcile ANTs-derived and FreeSurfer-derived segmentations of the cortical gray-matter of Mindboggle (Klein et al., 2017). Volume-based spatial normalization to standard space, MNI152NLin6Asym, was performed through nonlinear registration with antsRegistration (ANTs 2.2.0), using brain-extracted versions of both T1w reference and the T1w template. The following templates were selected for spatial normalization: FSL\u2019s MNI ICBM 152 non-linear 6th Generation Asymmetric Average Brain Stereotaxic Registration Model [Evans et al., 2012; TemplateFlow ID: MNI152NLin6Asym].

#### Functional data preprocessing

For each of the two BOLD runs found per subject (across all tasks and sessions), the following preprocessing was performed. First, a reference volume and its skull-stripped version were generated using a custom methodology of fMRIPrep. Head-motion parameters with respect to the BOLD reference (transformation matrices, and six corresponding rotation and translation parameters) are estimated before any spatiotemporal filtering using mcflirt (FSL 5.0.9, Jenkinson et al., 2002). BOLD runs were slice-time corrected using 3dTshift from AFNI 20160207 (Cox & Hyde, 1997). Susceptibility distortion correction (SDC) was omitted. The BOLD reference was then co-registered to the T1w reference using bbregister (FreeSurfer) which implements boundary-based registration (Greve & Fischl, 2009). Co-registration was configured with six degrees of freedom. The BOLD time-series (including slice-timing correction when applied) were resampled onto their original, native space by applying the transforms to correct for head-motion. These resampled BOLD time-series will be referred to as preprocessed BOLD in original space, or just preprocessed BOLD. The BOLD time-series were resampled into standard space, generating a preprocessed BOLD run in MNI152NLin6Asym space. Several confounding time-series were calculated based on the preprocessed BOLD: framewise displacement (FD), DVARS and three region-wise global signals. FD was computed using two formulations following Power (absolute sum of relative motions, Power et al., 2014) and Jenkinson (relative root mean square displacement between affines, Jenkinson et al., 2002). FD and DVARS are calculated for each functional run, both using their implementations in Nipype (following the definitions by Power et al. 2014). The three global signals are extracted within the CSF, the WM, and the whole-brain masks. Additionally, a set of physiological regressors were extracted to allow for component-based noise correction (CompCor, Behzadi et al., 2007). Principal components are estimated after high-pass filtering the preprocessed BOLD time-series (using a discrete cosine filter with 128s cut-off) for the two CompCor variants: temporal (tCompCor) and anatomical (aCompCor). tCompCor components are then calculated from the top 5% variable voxels within a mask covering the subcortical regions. This subcortical mask is obtained by heavily eroding the brain mask, which ensures it does not include cortical GM regions. For aCompCor, components are calculated within the intersection of the aforementioned mask and the union of CSF and WM masks calculated in T1w space, after their projection to the native space of each functional run (using the inverse BOLD-to-T1w transformation). Components are also calculated separately within the WM and CSF masks. For each CompCor decomposition, the k components with the largest singular values are retained, such that the retained components\u2019 time series are sufficient to explain 50 percent of variance across the nuisance mask (CSF, WM, combined, or temporal). The remaining components are dropped from consideration. The head-motion estimates calculated in the correction step were also placed within the corresponding confounds file. The confound time series derived from head motion estimates and global signals were expanded with the inclusion of temporal derivatives and quadratic terms for each (Satterthwaite et al., 2013). Frames that exceeded a threshold of 0.5 mm FD or 1.5 standardized DVARS were annotated as motion outliers. All resamplings were performed with a single interpolation step by composing all the pertinent transformations (i.e. head-motion transform matrices, susceptibility distortion correction when available, and co-registrations to anatomical and output spaces). Gridded (volumetric) resamplings were performed using antsApplyTransforms (ANTs), configured with Lanczos interpolation to minimize the smoothing effects of other kernels (Lanczos, 1964). Non-gridded (surface) resamplings were performed using mri_vol2surf (FreeSurfer).

### Hippocampal subfield ROI segmentation & registration

Hippocampal subfield regions of interest (ROIs) were segmented for each participant using the Automatic Segmentation of Hippocampal Subfields toolbox (ASHS, version 2.0.0; Hickling et al., 2024). Segmentation was performed using each participant’s high-resolution T2-weighted hippocampal slab (TSE) image together with the T1-weighted anatomical image, following standard ASHS procedures. The Hindy et al. (2016) young adult atlas was used for subfield labeling, yielding participant-specific labels for the left and right CA1, CA2/3, DG, subiculum (SUB), entorhinal cortex (ERC), perirhinal cortex (PRC), and parahippocampal cortex (PHC). All segmentations were visually inspected for anatomical accuracy aligning with quality control guidelines from Canada et al. (2024). These ASHS subfield labels were transformed into each subject-specific T1w functional analysis grid using a one-shot registration pipeline implemented using Advanced Normalization Tools (ANTs; Version 2.5.0; Avants et al., 2009).

Two rigid-body transformations matrices were calculated: (1) native TSE to native T1-weighted structural, and (2) native T1-weighted structural to the fMRIPrep-preprocessed T1w reference. These transforms were applied in a single resampling step, mapping each binarized ROI directly from native TSE space into the final analysis grid to minimize interpolation. The final target grid was defined by each participant’s first-level SPM contrast image in subject-specific T1w-aligned functional space. Nearest-neighbor interpolation was used to preserve the categorical integrity of ROI labels, and all transformed masks were binarized after resampling. Final ROIs were visually inspected to confirm alignment with the subject-specific T1w functional grid (see Figure 2 for example segmentation).

**Figure 2.**
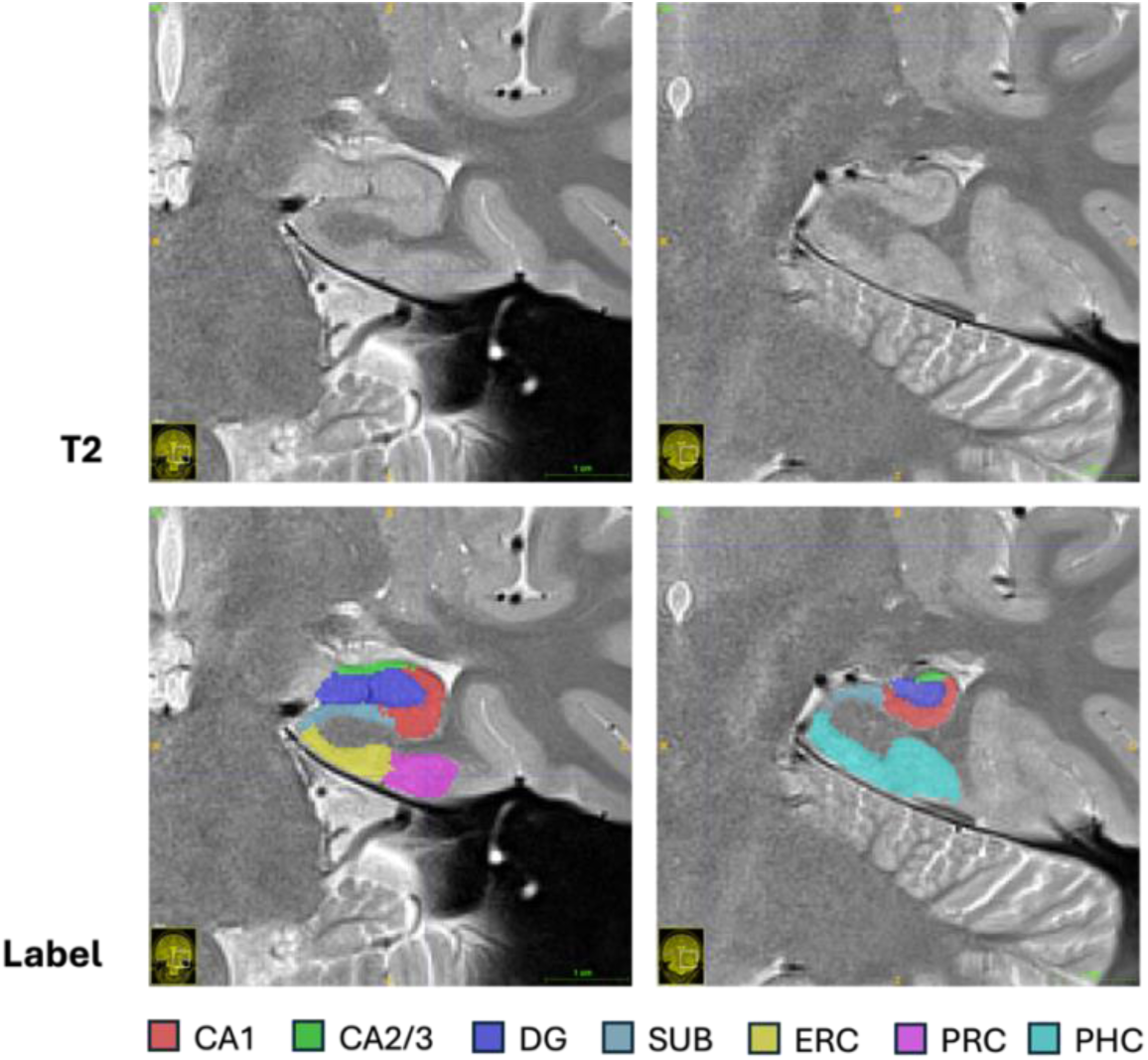
Example hippocampal subfield segmentation labels of one subject, in coronal view. High resolution T2 slab (top row) and segmented labels overlaid on top (bottom row).

### Behavioral analyses

Behavioral performance was assessed using two standard measures derived from the MDT: the Lure Discrimination Index (LDI) and corrected recognition memory (REC). The LDI provides a behavioral measure of mnemonic discrimination and was computed as the difference between the probability of giving a “similar” response to lures minus the probability of giving a “similar” response to foils. The REC provides a measure of traditional item recognition memory (i.e., the ability to correctly identify exact repetitions of studied items) and was calculated as the probability of giving a “same” response to targets minus the probability of giving a “same” response to foils.

To examine behavioral performance as a function of lag, LDI and REC were computed separately at each lag. Baseline response rates were calculated by averaging responses to foils within each subject to provide an estimate of response bias. Lag-specific LDI and REC scores were then computed by subtracting these baseline probabilities from lag-specific response probabilities. Lag effects on LDI and REC were tested using linear mixed-effects models using the lmer function from the lme4 package (Bates et al., 2015), with lag entered as a fixed effect and subject included as a random intercept. Post hoc pairwise comparisons across lag levels were conducted with model-estimated marginal means using the emmeans package (Lenth, 2017) with Benjamini–Hochberg correction for multiple comparisons.

### ROI analyses

ROI-based univariate analyses were conducted using the Statistical Parametric Mapping (SPM12; Ashburner et al., 2014) software in MATLAB R2023b (The MathWorks, Inc., 2023). Given our specific interest in hippocampal subfields, ROI analyses were conducted on unsmoothed functional data in subject-specific T1w-aligned functional space.

Trial-related activity was modeled at the first level using a general linear model (GLM) with stick functions at trial onset convolved with the canonical hemodynamic response function. Regressors of interest included all combinations of response type (same, similar, new) and trial type (first presentations, targets, lures, foils). For target and lure trials, separate regressors were created for each lag distance (10, 60, 140). First presentations and foils by response type were each modeled with a single regressor, as lag was not defined for these trial types. All remaining trials, including non-response trials, were modeled into a single regressor of no interest. Six motion parameters were included as nuisance regressors. Volumes exceeding a threshold of 0.5 mm FD or 1.5 standardized DVARS were identified as outliers during preprocessing. These volumes as well as the volumes immediately preceding and following each outlier, were modeled using single-volume nuisance (‘spike’) regressors of no interest.

The primary conditions of interest were “same” responses to targets (Hits) and “similar” responses to lures (LureCRs). Although successful mnemonic discrimination was the primary focus, Hits were also analyzed to provide a within-task comparison condition under the same lag manipulation, allowing assessment of whether lag-related neural effects were specific to mnemonic discrimination or reflected more general effects of temporal interference on recognition memory. Consistent with prior MDT studies, “new” responses to foils (FoilCRs) served as the baseline condition. Within each ROI, baseline-adjusted beta estimates were computed by subtracting the FoilCR beta weights from Hit or LureCR beta estimates (e.g., LureCR = LureCR – FoilCR).

To identify cortical regions associated with successful mnemonic discrimination beyond the MTL, a separate whole-brain univariate analysis was conducted. Whole-brain contrasts were estimated in MNI space using normalized functional images. This model included regressors corresponding to all combinations of response type and trial type, collapsed across lag to ensure that functional ROIs were defined independent of lag effects examined in subsequent analyses. All other modeling details were identical to the ROI analyses, except that spatially smoothed functional data (6mm FWHM Gaussian kernel) were used to increase sensitivity for detecting whole-brain effects.

Successful mnemonic discrimination was operationalized as greater activity for “similar” responses to lures (LureCRs) relative to “same” responses to lures (LureFAs), consistent with prior MDT work. Item recognition was operationalized using a separate contrast comparing “same” responses to targets (Hits) relative to LureFAs. LureFAs were used as the baseline due to low trial counts for target misses (“new” responses to targets). The resulting LureCR > LureFA and Hit > LureFA contrasts were thresholded at a voxelwise uncorrected threshold of *p* < .001 and cluster-corrected at *p* < .05, using 3dClustSim in AFNI Version 21.3.04 (Cox, 1996; Cox et al., 2017; Cox & Hyde, 1997). Significant clusters were used to define functional cortical ROIs (Tables 1 and 2).

**Table 1.** Significant clusters from whole-brain univariate analysis of Hit vs. LureFA. *k* = voxel size; H = hemisphere; L = left; R = right; *xyz* coordinates in MNI space.

| Region | H | $k$ | $t$ | *MNI coordinates* | | |
| --- | --- | --- | --- | --- | --- | --- |
| | | | | $x$ | $y$ | $z$ |
| Precuneus | R | 1829 | 7.13 | 5 | -73 | 35 |
| Ventral Striatum | L | 222 | 6.74 | -2 | 8 | -4 |
| Angular Gyrus | L | 476 | 6.30 | -49 | -68 | 32 |
| Medial Prefrontal Cortex | L | 925 | 5.71 | -7 | 54 | 1 |
| Temporoparietal Junction | R | 285 | 5.12 | 63 | -53 | 22 |

**Table 2.** Significant clusters from whole-brain univariate analysis of LureCR vs. LureFA. *k* = voxel size; H = hemisphere; L = left; R = right; *xyz* coordinates in MNI space.

| Region | H | $k$ | $t$ | *MNI coordinates* | | |
| --- | --- | --- | --- | --- | --- | --- |
| | | | | $x$ | $y$ | $z$ |
| Inferior Temporal Gyrus | L | 864 | 7.89 | -58 | -58 | -19 |
| Ventrolateral Prefrontal Cortex | L | 1141 | 7.19 | -42 | 53 | -14 |
| Superior Parietal Lobule | L | 768 | 6.35 | -30 | -72 | 49 |
| Cerebellum | R | 457 | 6.35 | 10 | -87 | -28 |
| Dorsolateral Prefrontal Cortex | L | 138 | 4.40 | -30 | 19 | 59 |

These functional cortical ROIs were analyzed alongside hippocampal subfield and MTL ROIs to investigate lag effects in mnemonic discrimination. Baseline-adjusted beta estimates were analyzed using linear mixed-effects models with the lmer function from the lme4 package (Bates et al., 2015). Separate models were fit for Hits and LureCRs within each ROI. In each model, beta estimates were modeled as a function of lag (e.g., 10, 60, 140), with a random intercept for subject. Fixed-effect significance was assessed using Type II Wald χ² tests via the car package’s Anova function (Fox & Weisberg, 2018). Planned pairwise contrasts were computed from model-estimated marginal means using the emmeans package (Lenth, 2017) and Benjamini-Hochberg corrected for multiple comparisons (Benjamini & Hochberg, 1995). Full list of cortical ROIs for Hits and LureCRs are provided in Tables 1 and 2, respectively.

### Exploratory whole-brain parametric modulation analyses

To identify regions beyond a priori and functionally defined ROIs whose activity differed by temporal interference level, we conducted an exploratory whole-brain parametric modulation analysis in SPM12. Using smoothed functional data, first-level GLMs included condition regressors for each trial-type-by-response combination. For target and lure trials, lag (10, 60, 140) was entered as a first-order (linear) parametric modulator to estimate lag-related changes in BOLD. At the group level, one sample t-tests were performed on subject-level lag-slope contrast images (e.g., Hit-by-lag; LureCR-by-lag). Resulting contrast maps were thresholded at *p* < .001 (uncorrected voxelwise) and cluster-corrected at *p* < .05, using 3dClustSim in AFNI Version 21.3.04 (Cox, 1996; Cox et al., 2017; Cox & Hyde, 1997).

## Results

### Behavioral performance

Figure 3A shows the proportions of “same,” “similar,” “new,” and no-responses to targets, lures, foils, and first presentations. We first assessed whether successful mnemonic discrimination and target recognition performances declined with increasing lag. Separate linear mixed-effects models were fit for REC and LDI, with lag as a fixed effect and subject as random intercept (Figures 3B and 3C). Consistent with our hypotheses, lag significantly affected both REC, χ²(2) = 72.55, *p* < .001, and LDI, χ²(2) = 162.08, *p* < .001. For both measures, performance was highest at lag 10 and declined from lag 10 to lag 60 and lag 140.

**Figure 3.**
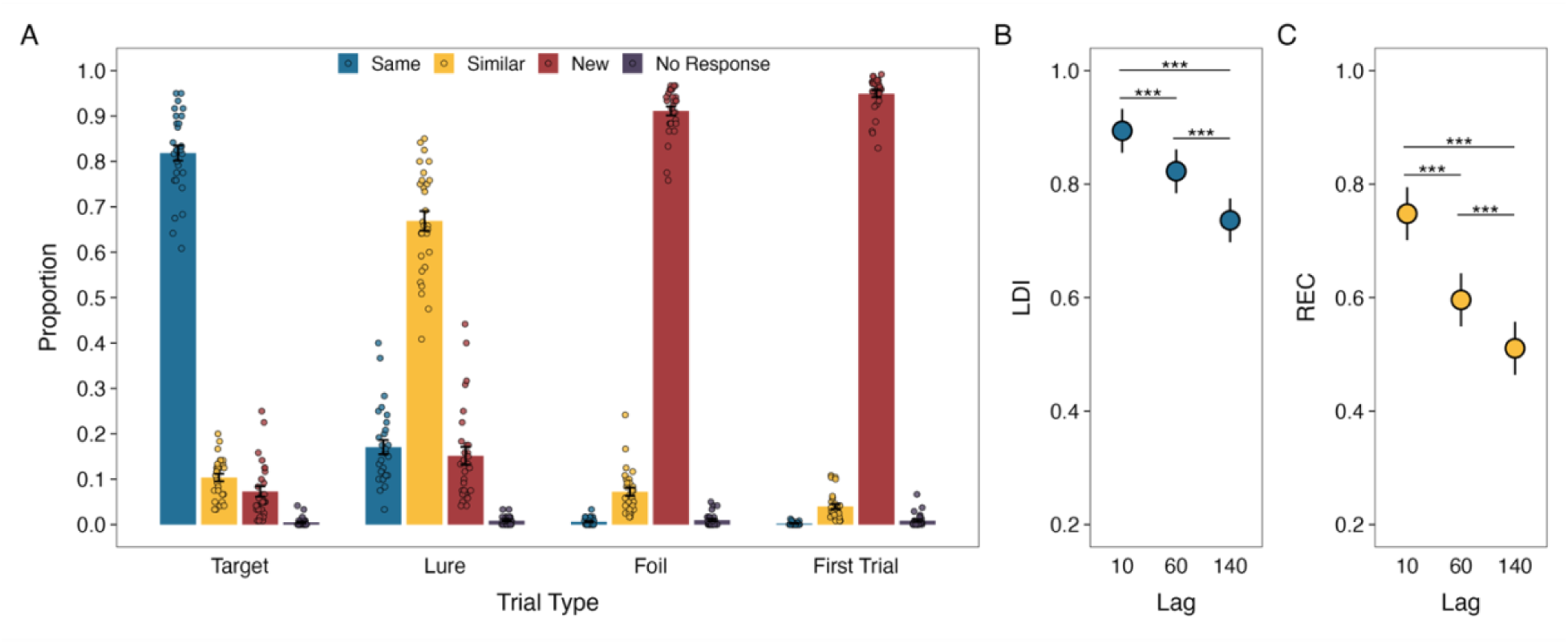
Proportion of responses conditioned on trial type (A), REC by lag (B) and LDI by lag (C). Points in (B) and (C) reflect model-estimated values from mixed-effects models; error bars represent 95% confidence intervals; \**p* < .05, \*\**p* < .01, \*\*\**p* < .001.

As follow-up analyses, to better characterize the response patterns underlying the lag effects, we examined lure trial responses using separate generalized linear mixed-effects models predicting probability of “same,” “similar,” and “new” responses as function of lag. All three models revealed significant main effects of lag, χ²(2) = 41.40, *p* < .001, χ²(2) = 153.22, *p* < .001, and χ²(2) = 100.36, *p* < .001, respectively. Pairwise comparisons revealed that the probability of “similar” responses to lures decreased with increasing lag: “similar” responses were more likely at lag 10 compared to both lag 60 (OR = 2.29, 95% CI [1.80, 2.90], *p* < .001) and lag 140 (OR = 3.36, 95% CI [2.65, 4.25], *p* < .001), and were also more likely at lag 60 compared to lag 140 (OR = 1.47, 95% CI [1.19, 1.81], *p* < .001). In contrast, the probability of “same” responses to lures was lower at lag 10 compared to lag 60 (OR = 0.51, 95% CI [0.38, 0.68], *p* < .001), and lag 140 (OR = 0.49, 95% CI [0.37, 0.66], *p* < .001) but did not significantly differ between lag 60 and lag 140 (OR = 0.97, 95% CI [0.76, 1.25], *p* = .794). The probability of “new” responses increased across lag conditions, with probability being lower lag 10 relative to lag 60 (OR = 0.48, 95% CI [0.35, 0.68], *p* < .001), and lag 140 (OR = 0.27, 95% CI [0.20, 0.37], *p* < .001), and lower at lag 60 relative to lag 140 (OR = 0.56, 95% CI [0.42, 0.73], *p* < .001).

### ROI analyses

To investigate lag effects on neural activity within hippocampal subfield, MTL and functionally defined cortical ROIs, beta estimates were analyzed using linear mixed-effects models. Separate models were fit for Hits and LureCRs within each ROI. In each model, lag was entered as a fixed effect, with subject included as a random intercept.

#### Hippocampal Subfield ROI analysis

For hippocampal subfields, neither the Hit nor LureCR models revealed a significant effect of lag, largest χ²(2) = 4.28, all *p*s ≥ .118, and largest χ²(2) = 5.76, *p*s ≥ .056, respectively (Figure 4).

**Figure 4.**
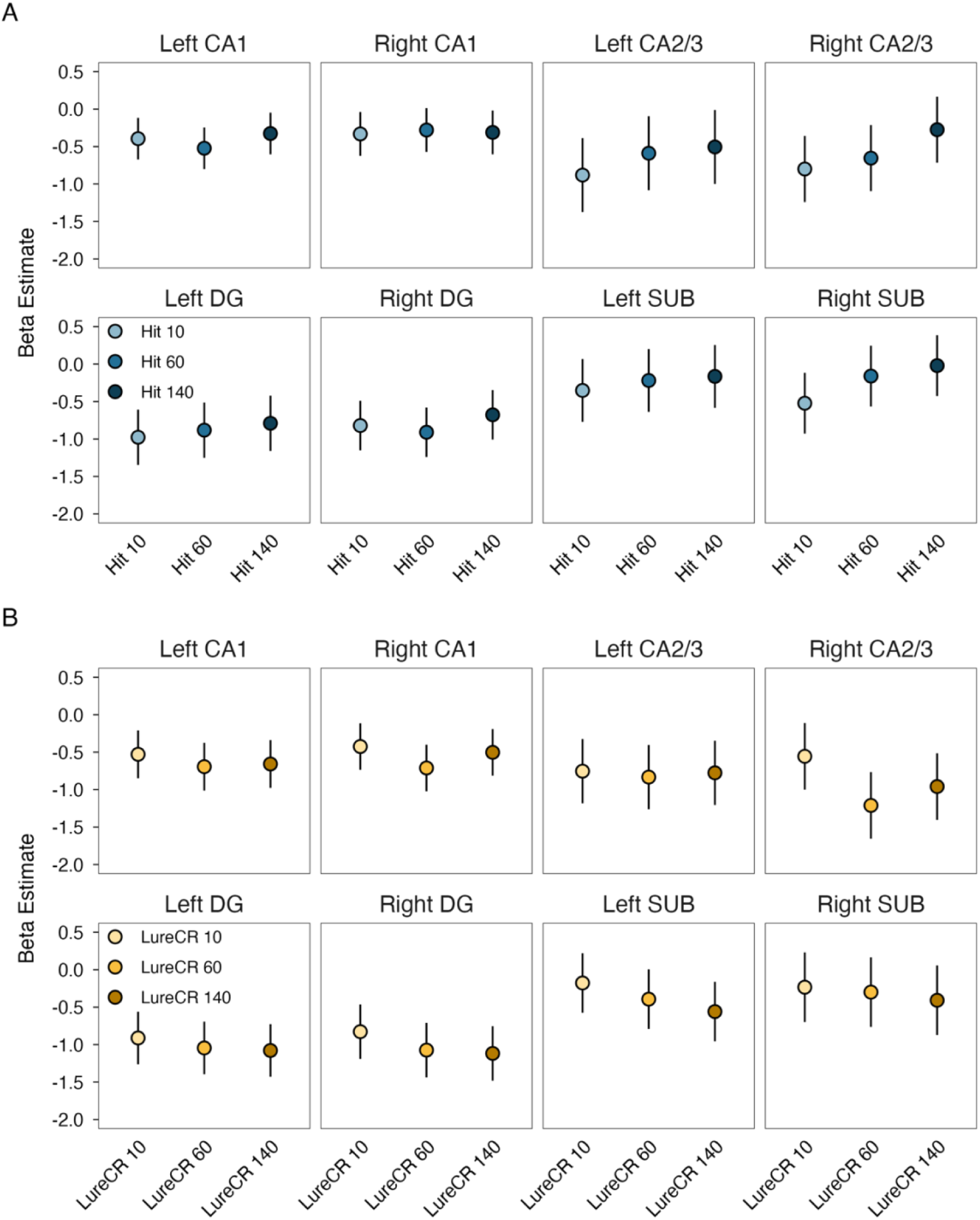
Beta estimates conditioned on lag for Hits (A) and LureCRs (B) across hippocampal subfields. Points reflect model estimated marginal means from mixed-effects models; error bars represent 95% confidence intervals; \**p* < .05, \*\**p* < .01, \*\*\**p* < .001.

#### MTL ROI analysis

Among MTL regions, significant effects of lag were observed for Hits in the left and right PHC, χ²(2) = 22.12, *p* < .001, χ²(2) = 21.186, *p* < .001, and the left and right PRC, χ²(2) = 8.53, *p* = .014, χ²(2) = 26.67, *p* < .001 (Figure 5A). Post-hoc comparisons revealed that, in both the left PHC and right PRC, beta estimates were lowest at lag 10 and increased with lag. In the right PHC, beta estimate for Hit10 was lower than lag 140 (*β* = −0.61, 95% CI [−0.94, −0.28], *p* < .001) and beta weight for lag 60 was also lower than lag 140 (*β* = −0.39, 95% CI [−0.72, −0.05], *p* = .009). However, activity did not differ between lags 10 and 60 (*β* = −0.23, 95% CI [−0.56, 0.11], *p* = .098). In the left PRC, beta estimates were significantly lower for at lag 10 compared to both lag 60 and lag 140, (*β* = −0.40, 95% CI [−0.76, −0.03], *p* = .029 and *β* = −0.35, 95% CI [−0.71, −0.02], *p* = .034), while activity did not differ between lag 60 and lag 140 (*β* = 0.05, 95% CI [−0.32, 0.41], *p* = .742).

**Figure 5.**
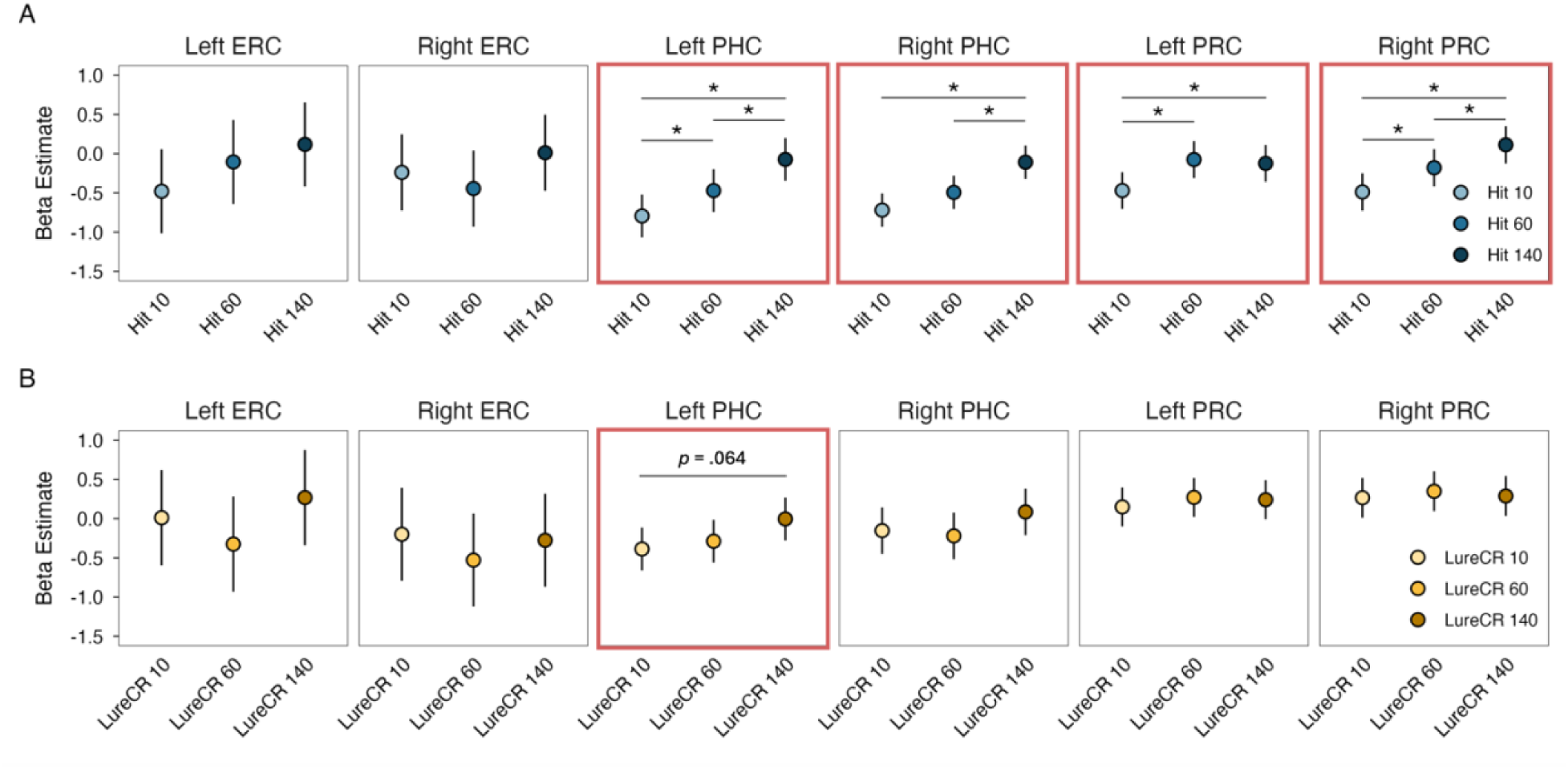
Beta estimates conditioned on lag for Hits (A) and LureCRs (B) across MTL regions. Points reflect model estimated marginal means from mixed-effects models; error bars represent 95% confidence intervals; \**p* < .05, \*\**p* < .01, \*\*\**p* < .001; red-border panels reflect significant omnibus effect of lag.

Within LureCRs, a significant effect of lag was observed in the left PHC, χ²(2) = 6.05, *p* = .049, but, none of the pairwise comparisons survived correction for multiple comparisons (smallest *p* = .064; Figure 5B).

#### Cortical ROI analysis

For functionally defined cortical ROIs, Beta estimates for Hits showed significant effects of lag in the medial prefrontal cortex (mPFC), χ²(2) = 9.69, *p* = .008, and the right temporoparietal junction (TPJ), χ²(2) = 20.056, *p* < .001 (Figure 6A). In the left mPFC, beta estimates at lag 140 were significantly greater than at lag 10 (*β* = 0.47, 95% CI [0.01, 0.93], *p* = .021) and lag 60 (*β* = 0.53, 95% CI [0.07, 0.98], *p* = .019). In contrast, in the right TPJ, beta-estimates were significantly higher at lag 10 than at lag 60 (*β* = 0.57, 95% CI [0.21, 0.93], *p* < .001), and 140 (*β* = 0.57, 95% CI [0.21, 0.93], *p* < .001).

**Figure 6.**
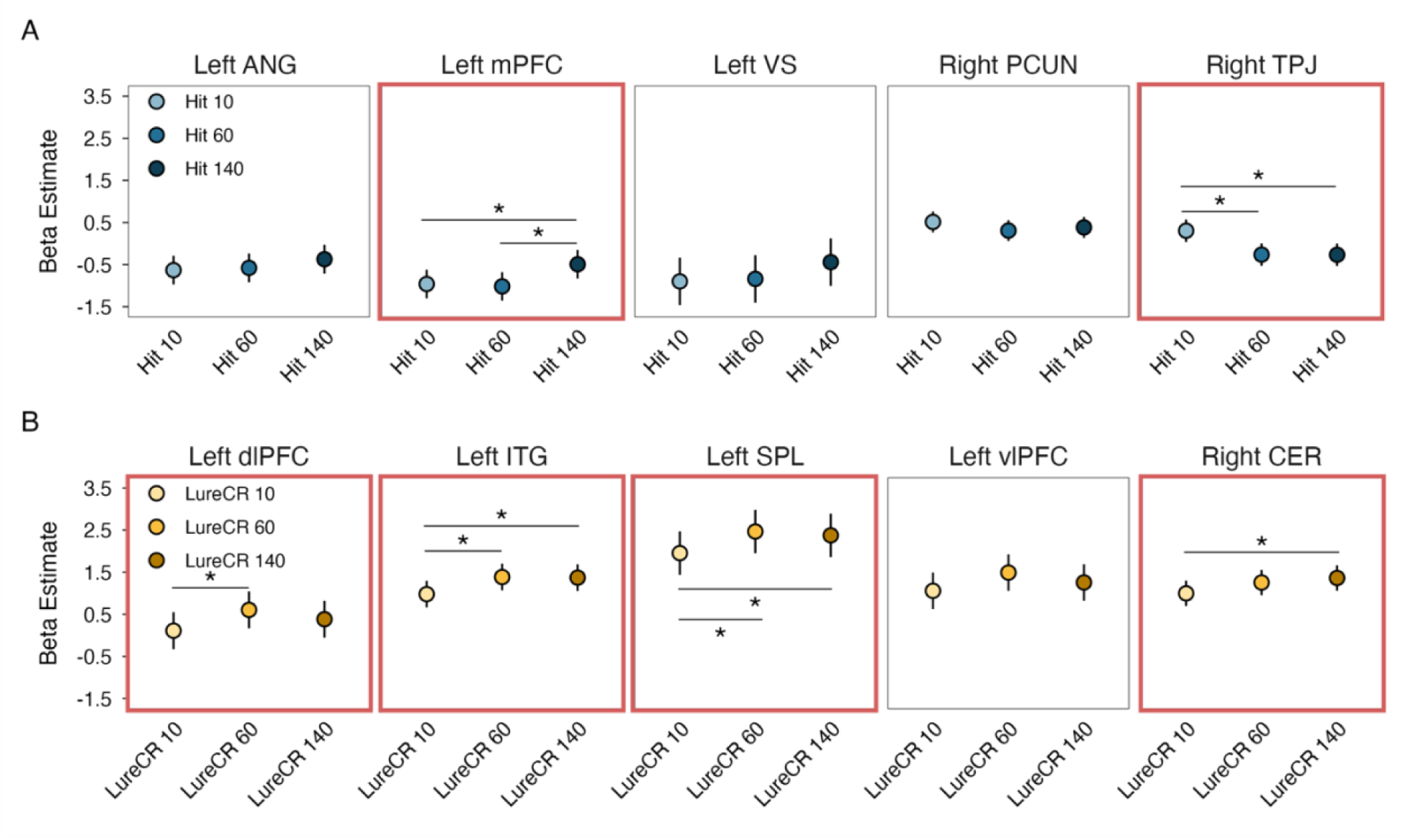
Beta estimates conditioned on lag for Hits (A) and LureCRs (B) across cortical regions. Points reflect model estimated marginal means from mixed-effects models; error bars represent 95% confidence intervals; \**p* < .05, \*\**p* < .01, \*\*\**p* < .001; red-border panels reflect significant omnibus effect of lag.

For LureCRs, significant effects of lag were observed in right cerebellum, χ²(2) = 7.623 *p* = .027, left superior parietal lobule (SPL), χ²(2) = 14.25, *p* < .001, left inferior temporal gyrus (ITG), χ²(2) = 17.67, *p* < .001, and left dorsolateral prefrontal cortex (dlPFC), χ²(2) = 7.47, *p* = .024 (Figure 6B). Post hoc comparisons revealed that in the left SPL, beta estimates were significantly lower at lag 10 compared to lag 60 (*β* = −0.51, 95% CI [−0.87, −0.15], *p* = .003) and lag 140 (*β* = −0.42, 95% CI [−0.78, −0.06], *p* = .008). A similar pattern was observed in the left ITG, with lower activity at lag 10 relative to lag 60 (*β* = −0.41, 95% CI [−0.68, −0.14], *p* = .001) and lag 140 (*β* = −0.39, 95% CI [−0.66, −0.12], *p* = .001). In the left dlPFC, beta estimate at lag 10 was significantly lower than at lag 60 (*β* = −0.50, 95% CI [−0.94, −0.05], *p* = .025). In the right cerebellum, beta estimate was significantly higher at lag 140, compared to lag 10 (*β* = 0.37, 95% CI [0.02, 0.71], *p* = .034).

### Exploratory whole-brain parametric modulation analysis

We conducted whole-brain parametric modulation analysis for Hits and LureCRs separately, and in conjunction, to identify brain regions whose activities increased with lag (see Table 3). Significant regions for Hits included the bilateral occipital pole, left vmPFC, left posterior cingulate gyrus, and left parahippocampal cortex. Significant clusters for LureCRs included the right superior frontal gyrus, bilateral lateral occipital cortex, right inferior frontal gyrus, right intraparietal sulcus, and left superior parietal lobule. Conjunction analyses of Hits and LureCRs revealed significant clusters in which activity scaled with lag within the visual cortex (see Table 3).

**Table 3.**
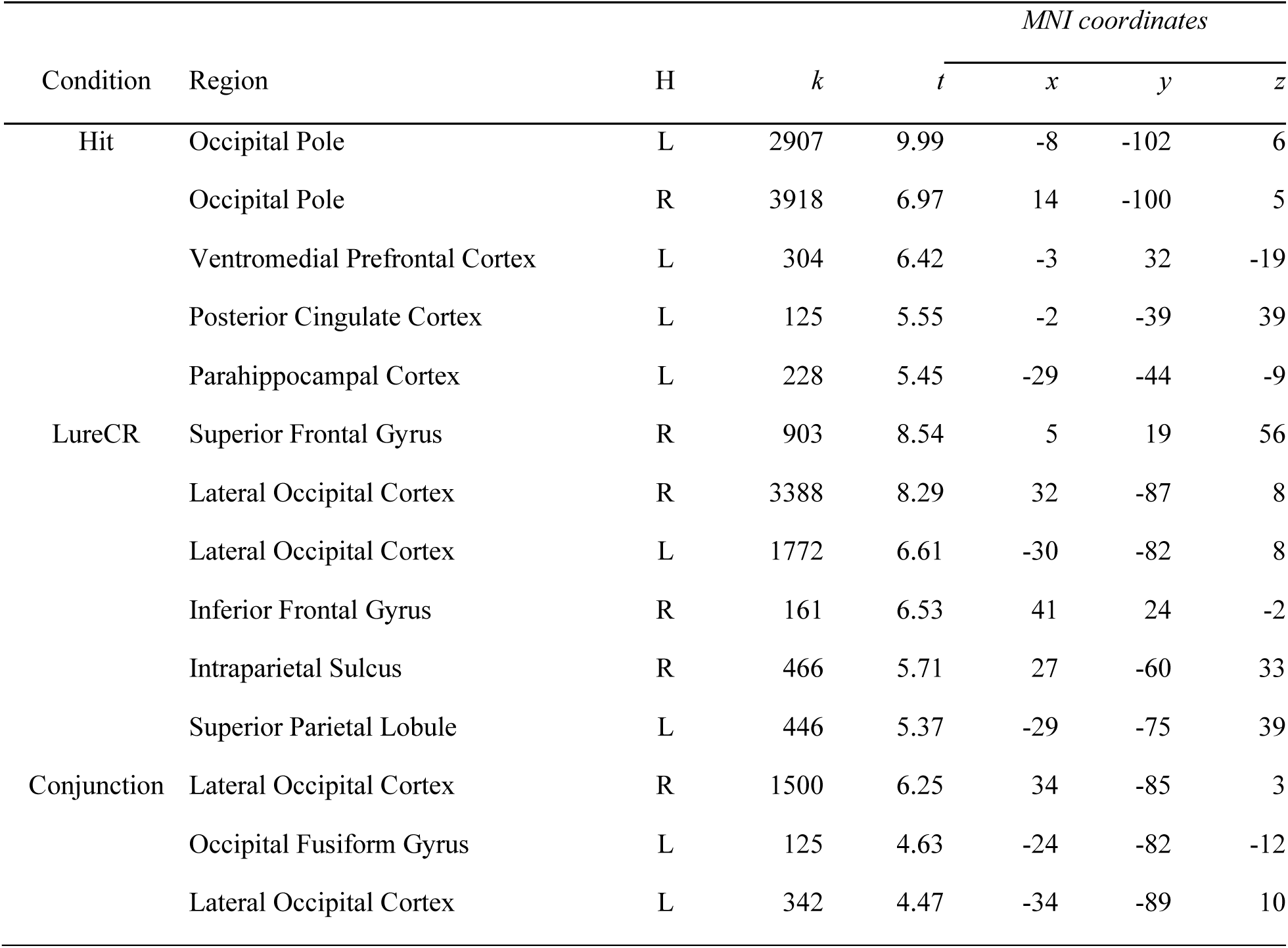
Significant clusters from whole-brain parametric modulation analysis for Hits, LureCRs and conjunction of Hits and LureCRs. *k* = voxel size; H = hemisphere; L = left; R = right; *xyz* coordinates in MNI space.

## Discussion

The current study investigated how temporal interference affects mnemonic discrimination and its neural correlates by systematically manipulating lag, or the number of trials that occurred between first presentations of images and subsequent target or lure presentations, in a continuous-recognition variant of the MDT. Consistent with interference-based models of memory, (McClelland et al., 1995, 2020; Norman & O’Reilly, 2003; Yassa & Reagh, 2013), increasing lag impaired both mnemonic discrimination (LDI) and target recognition (REC), indicating that greater temporal interference reduced overall memory performance. Partially contradicting our hypotheses, the neural effects of lag were absent in hippocampal subfields, but did emerge primarily within extrahippocampal MTL regions, including the PHC and PRC, as well as other cortical regions, including frontal and parietal cortices. Importantly, lag effects on neural activity differed across trial types such that neural activity patterns associated with successful mnemonic discrimination (LureCRs) were dissociable from those supporting successful target recognition (Hits). Converging with these findings, exploratory whole-brain parametric modulation analyses revealed largely distinct cortical regions whose activity scaled with lag for both LureCRs and Hits. Together, these findings highlight an important role of the medial temporal lobe and other cortical regions in resolving temporal interference to support mnemonic discrimination and recognition processes. Rather than reflecting computations restricted to hippocampal subfields alone, our results suggest that temporal interference modulates neural activity within broader MTL circuitry, including parahippocampal and perirhinal cortices, as well as frontal and parietal regions. These findings align with theoretical accounts proposing that pattern separation reflects distributed interactions between hippocampal and cortical systems (Amer & Davachi, 2023; Ritchey et al., 2015).

### Longer lags impair both mnemonic discrimination and target recognition

Our behavioral results revealed a robust lag effect, with memory performance declining for both mnemonic discrimination and target recognition. These findings align with prior studies using continuous-recognition MDTs (Ally et al., 2013; S. M. Stark et al., 2015), as well as paradigms employing separate study-test phase studies (Leal et al., 2019; Morales-Calva et al., 2026; Roberts et al., 2014), which have demonstrated that increasing temporal distance between encoding and retrieval impairs mnemonic discrimination. The decline in mnemonic discrimination performance (LDI) was primarily driven by a reduction in “similar” responses to lures, while both “same” and “new” responses increased with lag. Within continuous-recognition paradigms, lag effects are interpreted as reflecting increased temporal interference arising from the accumulation of intervening stimuli (Fox & Osth, 2023; Friedman, 1990; Yassa & Reagh, 2013). Specifically, as lag increases, participants encounter a greater number of items between an object’s initial presentation and its subsequent repetition or lure, resulting in heightened lure discrimination demands. In our study, with longer lags (i.e. heightened interference), participants were more likely to misclassify lures as exact repetitions (“same”), reflecting increased reliance on familiarity or pattern completion processes, as well as to classify lures as novel (“new”), reflecting retrieval failure or forgetting. The simultaneous increase in both misattribution errors and forgetting provides behavioral evidence that lag effects primarily reflect interference rather than mere time-based decay.

### Null lag effects in hippocampal subfields

Based on theoretical models of hippocampal pattern separation and prior empirical findings (Berron et al., 2016; Kirwan & Stark, 2007; Yassa & Stark, 2011), we hypothesized that the DG would be particularly sensitive to lag-related interference. Specifically, we predicted that DG activity would increase with lag, reflecting the increasing demands on DG-mediated pattern separation computations accruing with the increasing temporal interference. However, despite robust behavioral lag effects, none of the hippocampal subfields showed lag-dependent modulation of neural activity during successful mnemonic discrimination.

Several factors may account for this null effect. Successful mnemonic discrimination was observed across all lag conditions, suggesting that hippocampal pattern separation mechanisms were likely engaged throughout the task. Under this assumption, hippocampal activity may have remained relatively stable once temporal interference exceeded a threshold sufficient to recruit pattern separation processes. Alternatively, although lag reliably influenced behavioral performance, the magnitude of interference differences between lag conditions may not have been sufficient to elicit detectable differences in mean BOLD amplitude within hippocampal subfields. The fact that hippocampal subfield activity did not differ across lag conditions for Hits further supports this account. Additionally, with regard to target/lure relatedness, all lure trials were drawn from lure similarity level 4 (S. M. Stark et al., 2019), corresponding to relatively low perceptual overlap between first presentations and their later lures (i.e., less similar). Although this design choice enabled stricter experimental control to isolate temporal interference from lure similarity effects, reduced overall similarity between studied items and lures may have attenuated demands on hippocampal pattern separation mechanisms.

Another contributing factor may be the continuous-recognition structure of the task in combination with its intentional memory decision demands. Unlike separate study-test paradigms, our MDT required participants to engage in ongoing memory decisions throughout the entire experiment. Although prior work indicates that continuous and study-test MDT variants produce comparable behavioral outcomes (C. E. L. Stark et al., 2023; S. M. Stark et al., 2015, 2019), these paradigms may differ in their ability to tease apart specific neural signatures of behavioral pattern separation and other mnemonic processes. Continuous memory judgments on the present task may have induced a sustained retrieval state, potentially elevating baseline hippocampal engagement for both pattern separation and pattern completion across lag conditions. Under such circumstances, lag-related differences may have produced weaker effects on the mean hippocampal BOLD activity observed, even when behavioral effects were significant. Consistent with this interpretation, many studies demonstrating pattern separation activity in the DG have employed incidental paradigms, in which participants are not required to make explicit memory judgments about lure stimuli (Bakker et al., 2008; Lacy et al., 2011), whereas intentional memory tasks have been associated with more complex hippocampal activity patterns (Kirwan & Stark, 2007; Nash et al., 2021).

### Lag effects in extrahippocampal MTL regions

In contrast to hippocampal subfields, activity in other MTL regions, specifically the PRC and PHC, showed robust lag effects. Contrary to our hypotheses within the PRC, neural activity for target recognition (Hits), but not mnemonic discrimination (LureCRs), varied by lag.

Specifically, neural activity for Hits increased monotonically with lag. This pattern is consistent with models of recognition memory proposing an important role of PRC in item-based recognition and familiarity-related processes (Brown & Aggleton, 2001; Diana et al., 2007; Ranganath & Ritchey, 2012). According to these models, successful target recognition does not necessarily require retrieval of detailed episodic information but may instead rely on assessments of global memory strength or familiarity signals. Within this account, lag-related increases in PRC activity likely reflect greater reliance on familiarity-based recognition under increased temporal interference. As lag increases, intervening items introduce greater mnemonic interference, reducing the accessibility of target-specific episodic information. Accordingly, recognition decisions may increasingly depend on familiarity signals rather than precise retrieval of prior representations. Under this account, lag-related PRC modulation would be expected for Hits, which can be accurately identified based on familiarity even when episodic specific is reduced, but not for LureCRs, which require more detailed discrimination among overlapping representation. Additionally, prior work has shown that PRC activity is uniquely affected by object-based interference, suggesting that PRC activity buffers against the interfering effects of competing object information on recognition memory (Watson & Lee, 2013). The PRC has also been shown to support reinstatement of item-level representations during retrieval (Tompary et al., 2016), a process frequently associated with pattern completion mechanisms that contribute to successful target recognition.

While lag-dependent activity in the PRC was only observed for Hits, activity within the left PHC varied as a function of lag for both Hits and LureCRs. Unlike the PRC, the PHC is more frequently associated with recollection processes (Diana et al., 2007, 2013; Li et al., 2016; Sommer et al., 2005). Dual-process models propose that recollection involves retrieval of specific contextual or featural details from a prior event or information, whereas familiarity reflects a general sense that an item was previously encountered in the absence of detailed retrieval (Yonelinas, 2002). From this perspective, increasing lag, and the accompanying rise in temporal interference, likely amplified demands on recollection memory processes necessary to resolve competition among similar representations. Such processes would be critical not only for discriminating lures, but also for accurately endorsing previously studied targets under heightened interference. Converging with these findings, a prior high-resolution whole-brain fMRI study reported left PHC activity during successful mnemonic discrimination, although effects were not observed in other MTL regions (Nash et al., 2021), suggesting that PHC may be particularly sensitive to interference-related demands during discrimination of similar events.

### Lag effects in cortical regions

To investigate lag effects in cortical regions, ROIs were defined separately for target recognition (Hits; Hit > LureFA) and mnemonic discrimination (LureCR > LureFA), based on the assumption that these processes engage partially distinct neural mechanisms. Lag effects for Hits were observed in the left medial prefrontal cortex (mPFC/ACC/vmPFC) and right temporoparietal junction (TPJ). Within the left mPFC, Hit activity was greatest at the longest lag (140 trials), while activity at lag 10 and lag 60 did not differ. A large body of work has implicated mPFC regions including the medial superior frontal gyrus and anterior cingulate cortex, in top-down monitoring and response selection under conditions of uncertainty or increased difficulty in retrieval monitoring (Achim & Lepage, 2005; Cruse & Wilding, 2009; Euston et al., 2012; Ridderinkhof et al., 2004). In the current study, the longer lags likely reduced mnemonic specificity for target trials, shifting retrieval reliance toward familiarity-based signals. Under such conditions, increased recruitment of mPFC at longer lags may likely reflect a greater engagement of decision-related and retrieval-monitoring processes required to evaluate these weakened memory signals.

In contrast to the pattern of activity observed in the mPFC, the right TPJ activity for Hits showed activation greatest at the shortest lag (10), while activation at lag 60 and 140 did not differ. Within attentional accounts of parietal contributions to memory, TPJ activity is posited to reflect bottom-up capture of attention by retrieved information rather than active retrieval effort (Cabeza et al., 2008; Corbetta & Shulman, 2002; Wu et al., 2015). Under this framework, stronger or more vivid mnemonic signals, as expected at shortest lag (i.e., lag 10), are more likely to engage TPJ-mediated attentional processes, which is consistent with the elevated TPJ activity at the shortest lag in our study.

Cortical ROI analyses for LureCRs revealed significant lag effects within the left superior parietal lobe (SPL) and left dorsolateral prefrontal cortex (dlPFC). Both regions are central components of the frontoparietal control network implicated in attentional selection and interference resolution (Amer & Davachi, 2023; Harding et al., 2015; Spreng et al., 2010). Interestingly, our results show that activity in these regions increased from lag 10 to lag 60, but not from lag 60 to lag 140. This pattern suggests that at highest levels of temporal interference (i.e lag 140 trials), frontoparietal recruitment reaches a functional asymptote. That is, once interference exceeds a threshold sufficient to recruit attentional control and interference resolution processes, additional increases in lag may no longer induce proportional increase in neural activity. This pattern is consistent with models proposing that cognitive control systems operate in a demand-dependent but capacity-limited way, where neural recruitment increases with interference up to a certain capacity limit (Duncan, 2010; Miller & Cohen, 2001; Steffener et al., 2020).

The results of the whole-brain parametric modulation analyses further emphasize the role of frontoparietal control regions in mnemonic discrimination. During LureCR trials, neural activity increased linearly with lag across regions including the SFG, IFG, IPS, and SPL, all of which are regions in the frontoparietal control network (Dixon et al., 2018). Notably, these regions were absent for lag effects during target recognition, consistent with the view that target recognition, relative to mnemonic discrimination, requires less demands for interference resolution and detailed recollection processes (Yassa & Stark, 2011). Collectively, the cortical-ROI and whole-brain parametric modulation findings converge to suggest that increasing lag modulates cortical regions implicated in resolving interference. Notably, these robust lag effects in cortical regions may explain the absence of lag-related differences within hippocampal subfields. If increasing temporal interference was resolved, at least in part, by sensory/perceptual and cognitive control cortical regions, additional recruitment of hippocampal subfields may not have been necessary to support successful mnemonic discrimination across lag conditions.

In addition to our hypothesized regions, activity in the left inferior temporal gyrus (ITG) and right cerebellum were also associated with successful mnemonic discrimination, and were also subsequently modulated by lag. The inferior temporal gyrus is a key component of the ventral visual stream and plays a central role in object recognition (Conway, 2018; Majaj et al., 2015). Relatedly, a prior study (Doxey & Kirwan, 2015) found that the structural integrity of the inferior temporal gyrus was correlated with mnemonic discrimination performance. Engagement of this region during LureCR trials may reflect the fact that the current study employed an object-based MDT. With respect to cerebellar activity, several prior studies have implicated an important role of the cerebellum in episodic memory recall (Almeida et al., 2023; Andreasen et al., 1999; Fliessbach et al., 2007; Geissmann et al., 2023; Sweatman et al., 2023). Studies have also identified cerebellar networks that are functionally connected to the hippocampus during episodic memory tasks, one during free recall (Geissmann et al., 2023), and another during mnemonic discrimination (Paleja et al., 2014). Our findings extend this literature by suggesting that cerebellar engagement may scale with interference-related demands during mnemonic discrimination, further specifying its contribution to episodic memory processes. Together, these results indicate that mnemonic discrimination under conditions of increasing temporal interference engages a broader distributed network that extends beyond canonical MTL and frontoparietal regions. Importantly, these regions do not merely support mnemonic discrimination in general, but appear to be specifically engaged as interference demands increase, emphasizing that the resolution of temporal interference is implemented across distributed cortical regions. Future research will be necessary to clarify the precise contributions of these regions to interference resolution and how they interact with MTL circuitry to support successful mnemonic discrimination.

## Conclusions

In sum, the current study demonstrates that increasing temporal interference impairs both mnemonic discrimination and target recognition in a continuous-recognition MDT, and that the neural consequences of lag are robust in extrahippocampal MTL and cortical regions. Although hippocampal subfield-level activity did not vary by lag, activity in PRC and PHC showed clear lag-related modulation, with dissociable patterns for mnemonic discrimination and target recognition. Extending beyond the MTL, cortical lag effects were process-specific: mPFC and TPJ tracked target recognition whereas frontoparietal control regions were selectively modulated by lag during successful mnemonic discrimination. Together, these findings converge with cortico-hippocampal accounts of pattern separation, such as the CHiPS framework, suggesting that increasing temporal interference increases recruitment of both MTL and cortical regions that support interference resolution (Amer & Davachi, 2023; Ritchey et al., 2015).

Future work that differently manipulates interference demands, for example, by varying perceptual overlap between studied items and lures or by parametrically increasing lure similarity, will be critical for determining the conditions under which temporal interference modulates hippocampal subfield computations versus when interference can be resolved by distributed cortical control and perceptual regions that project to the hippocampus.

## Acknowledgments

The authors acknowledge use of the Social, Life, and Engineering Sciences Imaging Center (SLEIC), Pennsylvania State University, University Park, PA (RRID:SCR_014922). We acknowledge the use of the Human MRI Facility for MRI data collection, experimental consultation, and technical support.

